# RNAcontacts, a pipeline for predicting contacts from RNA proximity ligation assays

**DOI:** 10.1101/2022.10.06.511089

**Authors:** Sergey Margasyuk, Mariia Vlasenok, Guo Li, Changchang Cao, Dmitri D. Pervouchine

## Abstract

**Background:** High-throughput RNA proximity ligation assays are molecular methods that simultaneously analyze spatial proximity of many RNAs in living cells. Their principle is based on cross-linking, fragmentation, and consequent religation of RNAs followed by high-throughput sequencing. The generated fragments have two distinct types of splits, one resulting from pre-mRNA splicing, and the other resulting from ligating spatially close RNA strands.

**Findings:** Here, we present *RNAcontacts*, a universal pipeline for detecting RNA-RNA contacts in high-throughput RNA proximity ligation assays. It circumvents the inherent problem of mapping sequences with two distinct split types using a two-pass alignment, in which splice junctions are inferred from a control RNA-seq experiment on the first pass and then provided to the aligner on the second pass as *bona fide* introns. This approach allows for a more sensitive detection of RNA contacts and has higher specificity with respect to splice junctions that are present in the biological sample in comparison to previously developed methods. *RNAcontacts* extracts contacts, clusters their ligation points, computes the read support, and generates tracks for the visualization through the UCSC Genome Browser. It is implemented in a reproducible and scalable workflow management system *Snakemake* that allows fast and uniform processing of multiple datasets.

**Conclusions:** *RNAcontacts* represents a generic pipeline for the detection of RNA contacts that can be used with any proximity ligation method as long as one of the interacting partners is RNA.

*RNAcontacts* is available via github at https://github.com/smargasyuk/RNAcontacts/

## 1 Introduction

Rapid evolution of high-throughput sequencing technology enabled *in vivo* identification of spatial contacts between nucleic acids, including DNA contacts in 3D chromatin structure [1, 2, 3], functional enhancer-promoter interactions [4, 5], and chromatin-associated RNA-DNA contacts [6, 7]. These methods rest on the basic principle of digestion of nucleic acids crosslinked in macromolecular complexes and subsequent stochastic religation, which occurs predominantly between spatially proximal molecules. Deep sequencing of the resulting chimeric fragments produces hundreds of millions of reads encoding sequence signatures of the interacting loci.

A number of recently developed methods applied RNA proximity ligation to trace back RNA-RNA interactions *in vivo* and *in vitro* (see [8] and [9] for review). Some of them, such as PARIS [10], LIGR-seq [11], SPLASH [12], and COMRADES [13] use psoralen derivatives to induce reversible crosslinking between RNA duplexes to assess pairwise structural interactions *in vivo* with high sensitivity and specificity. In the RIC-seq protocol, the RNA strands are crosslinked through an RNA-binding protein (RBP) [14]. This approach not only recapitulates RNA secondary and tertiary structures but also facilitates the generation of three-dimensional maps of the interacting RNA-RBP complexes. In all these cases, the interactions are encoded within chimeric RNA sequences obtained via digestion and religation.

However, unlike DNA-DNA interactions, which manifest themselves in the sequencing data as split reads that align to a pair of genomic loci, RNA-RNA interactions may produce more sophisticated split alignments because pre-mRNAs are spliced. In particular, the crosslinked fragments may contain both exon-exon and proximity ligation junctions, thus producing short reads with both the canonical intronic GT/AG splits resulting from splicing, as well as non-GT/AG splits resulting from religation (Figure 1A). Accurate mapping of such reads is challenging because most short read alignment tools can deal only with one type of splits. With just one split model, the aligner would have a tradeoff between increasing the penalty for non-GT/AG splits to correctly identify introns at the expense of missing RNA-RNA contacts, or relaxing the GT/AG requirement to correctly detect RNA-RNA contacts while having incorrect intron mappings. Therefore, it is of interest to develop a computational method that maps short reads with such distinct split types. The current work describes a computational pipeline that achieves this goal without developing specialized alignment software.

**Figure 1:**
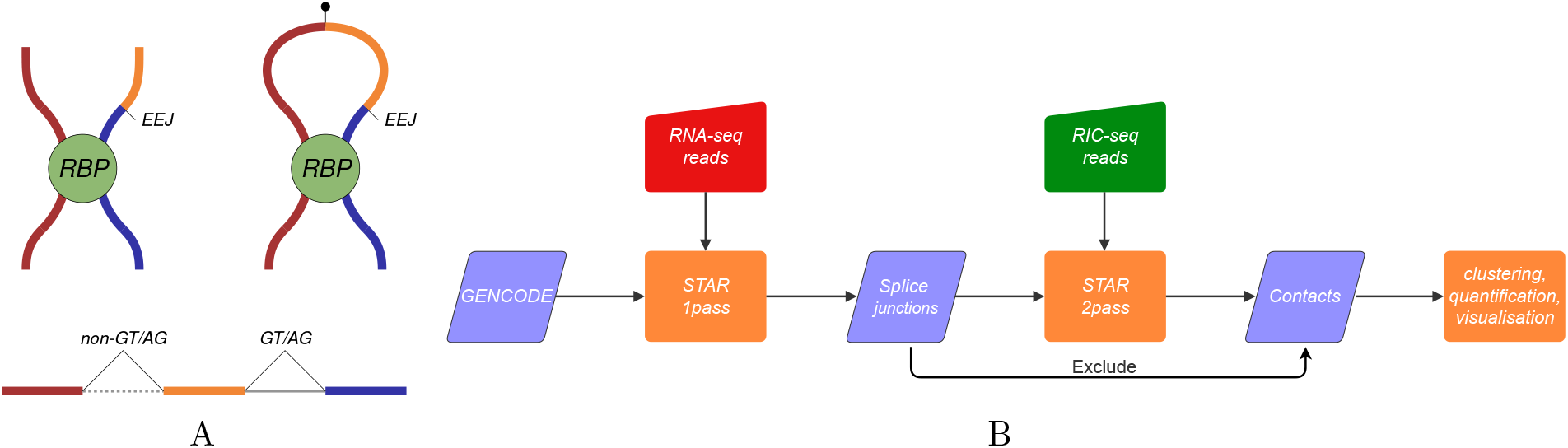
(A) In RIC-seq protocol [14], unspliced RNA strand can be crosslinked through an RNA-binding protein (RBP) to a strand with exon-exon junctions (EEJ). The sequence formed by proximity ligation aligns to the reference genome with a non-GT/AG split reflecting the ligation point and a canonical GT/AG split resulting from splicing. (B) The *RNAcontacts* pipeline. On the first pass, short reads from the control RNA-seq experiment are aligned to the reference genome to identify the expressed splice junctions. The latter are used on the second pass as *bona fide* introns when aligning proximity ligation data to detect ligation junctions, which encode RNA-RNA contacts.

## 2 Results

### 2.1 The RNAcontacts pipeline

Here, we present *RNAcontacts*, a computational pipeline for the analysis of RNA proximity ligation data, which circumvents the problem of multiple split types by aligning short reads in a two-pass mode (Figure 1B). The workflow is based on STAR aligner [15]. On the first pass, *RNAcontacts* aligns the sample-matched set of RNA-seq data in the paired-end mode to identify splice junctions that are expressed in the given biological sample using strict penalty for non-GT/AG splits. Here, RNA-seq represents a control experiment that doesn’t contain fragments resulting from proximity ligation. On the second pass, the reads generated in an RNA proximity ligation experiment are aligned using relaxed penalty for non-GT/AG splits, while providing splice junctions that were identified on the first pass as *bona fide* introns so that the aligner will preferentially make a split at the coordinates from the provided list. Since RNA proximity ligation data may contain chimeric junctions at arbitrary genomic distances or *in trans*, the alignment on the second pass is performed in single-end mode. All split alignments obtained on the second pass are parsed to extract RNA-RNA contacts and exclude splice junctions obtained on the first pass.

Spliced alignment programs usually generate two separate output files corresponding to colinear and non-co-linear splits. In particular, STAR aligner reports co-linear splits (same strand, same chromosome, and forward split orientation) within the standard SAM/BAM output, while non-co-linear splits are placed in the chimeric output [15]. *RNAcontacts* extracts the coordinates of neo-junctions, i.e., co-linear splits that were found on the second pass from the SAM/BAM output and combines them with chimeric splits obtained from the chimeric output. Of note, not only *trans* but also *cis* contacts may be encoded within both neo-junctions and chimeric splits. The combined output of the second pass consists of neo-junctions and chimeric junctions, which are jointly referred to as ligation junctions, and their constituent split positions are referred to as ligation points.

In application to RIC-seq experiment in the HeLa cell line [14], *RNAcontacts* mapped 94.3% of ∼224 million short reads from two bioreplicates, where 72.0% of the mapped reads were aligned uniquely (see Table S1 for complete mapping statistics). At that, 18.5% uniquely mapped reads contained at least one ligation junction, as compared to 3.5%, 2%, and 0.5% previously reported for LIGR-seq, PARIS, and SPLASH, respectively [16]. However, spliced alignment programs differ in their base-wise mapping accuracy and decisions on gap placement [17]. In application to RIC-seq protocol, for instance, gap variability can arise even when mapping read mates that overlap the same ligation point because reading the same sequence from one or the other strand may produce slightly different split coordinates due to the lack of consensus sequences that characterize the split (Figure S1). Furthermore, different copies of the same RNA are digested and re-ligated stochastically, thus resulting in even larger variability. Considering this technical and biological variation, we expect to observe clusters of ligation points rather than well-defined junctions as in spliceosomal GT/AG introns.

Indeed, the distribution of distances between two consecutive ligation points has a rapidly decaying tail, with approximately 50% of distances being below 9 nts and 90% of distances being below 21 nts (Figure S2A). We chose to cluster the ligation points using single linkage clustering with distance cutoffs (*δ*) of 10 nts and 20 nts. A contact was defined as a pair of clusters, with the number of supporting reads equal to the sum of read counts corresponding to ligation junctions (Table 1). For each *δ*, we subdivided the contacts into three groups: the intragene (both ends of a contact belong to an annotated gene), contacts *in cis* (on the same chromosome, but not in the same gene), and contacts *in trans* (on different chromosomes). The number of contacts (*n*), the cluster length (*s*), the distance between contacting clusters (*d*, which is defined only for intragene and *cis* contacts), and the number of supporting reads (*r*) were only marginally different for the two values of *δ*. On average, we detected 30% more intragene contacts than contacts *in cis*, and more than twofold enrichment of contacts *in trans* with respect to the other two groups. For *δ* = 10, most clusters had length 10 nts (Figure S3) indicating that they consist of only one individual ligation point surrounded by 5-nt-flanks in both directions.

**Table 1:**
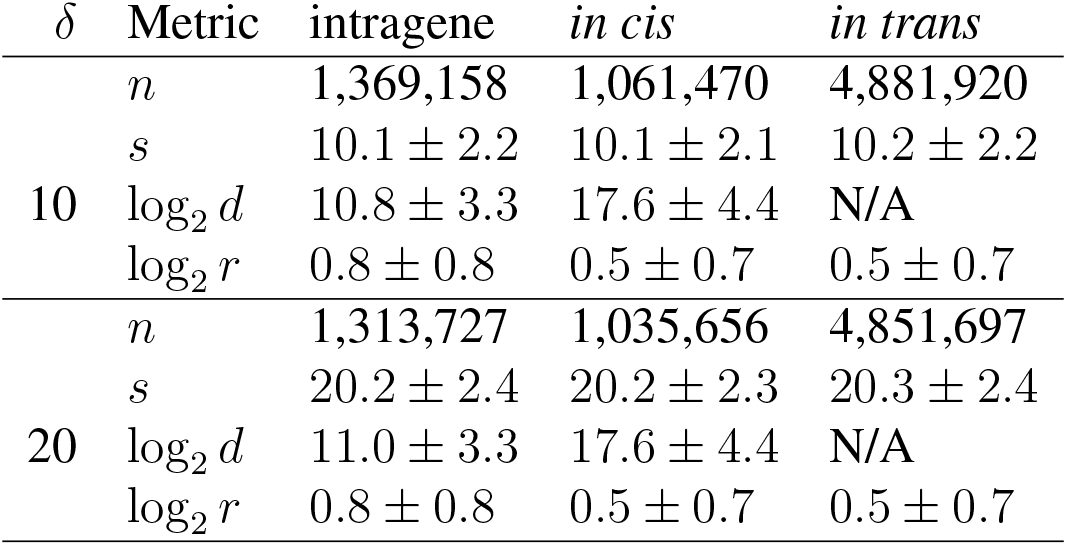
The characterization of clusters of RNA-RNA contacts. Clustering distance, *δ*. The number of contacts, *n*. Cluster lengths, *s*. Distance between contacting clusters, *d*. The number of reads supporting the contact, *r*. The numbers represent mean ± standard deviation.

| $\delta$ | Metric | intragene | *in cis* | *in trans* |
| --- | --- | --- | --- | --- |
| 10 | $n$ | 1,369,158 | 1,061,470 | 4,881,920 |
| | $s$ | $10.1 \pm 2.2$ | $10.1 \pm 2.1$ | $10.2 \pm 2.2$ |
| | $\log_2 d$ | $10.8 \pm 3.3$ | $17.6 \pm 4.4$ | N/A |
| | $\log_2 r$ | $0.8 \pm 0.8$ | $0.5 \pm 0.7$ | $0.5 \pm 0.7$ |
| 20 | $n$ | 1,313,727 | 1,035,656 | 4,851,697 |
| | $s$ | $20.2 \pm 2.4$ | $20.2 \pm 2.3$ | $20.3 \pm 2.4$ |
| | $\log_2 d$ | $11.0 \pm 3.3$ | $17.6 \pm 4.4$ | N/A |
| | $\log_2 r$ | $0.8 \pm 0.8$ | $0.5 \pm 0.7$ | $0.5 \pm 0.7$ |

The distances between contacting clusters were distributed differently for neo-junctions and chimeric splits, both in terms of the number of contacts and also when weighted by the number of supporting reads (Figure S4). Remarkably, the distributions have two modes, with the first mode at *d* ≃ 1, 000 corresponding to the intragene contacts encoded by both neo-junctions and chimeric splits. Chimeric reads may encode intragene contacts if the split is in backward orientation, as in circular RNAs [18]. The second mode for neo-junctions was due to the condition *d* ≤ 250, 000 that is imposed by STAR aligner on co-linear splits, however longer contacts *in cis* were captured by the second mode of the chimeric distribution. At that, most contacts *in cis* and *in trans* were supported by only one read, while most intragene contacts were supported by two reads (Figure S5). Therefore, the read support in individual RNA proximity ligation assays is generally quite sparse even after merging contacts into clusters.

### 2.2 RNAcontacts has better sensitivity than RICpipe

To compare the performance of *RNAcontacts* to that of *RICpipe*, a pipeline that was designed originally to analyze RIC-seq data, we first analyzed ligation junctions of 50 nts or longer that were located on the same chromosome and matched their exact genomic positions obtained by the two pipelines. We excluded junctions in reads mapped to rRNA from *RNAcontacts* results for this analysis because *RICpipe* removes rRNA reads [14]. Approximately 40% (respectively, 45%) of ligation junctions identified by *RNAcontacts* (respectively, *RICpipe*) had exactly the same coordinates as ligation junctions identified by the other pipeline likely indicating the difference in spliced alignment programs used (Figure 2A). However, in terms of the number of short reads supporting the identified ligation junctions *RNAcontacts* was able to align more reads as compared to *RICpipe*, indicating approximately 40% increase in sensitivity (Figure 2B). In carrying out the comparison using 100-nts windows, i.e., without exact coordinate matching, we observed that the results of the two pipelines are largely concordant, as also evidenced by highly similar contact maps with slightly more contacts for *RNAcontacts* as compared to RIC-pipe (Figure 2C).

**Figure 2:**
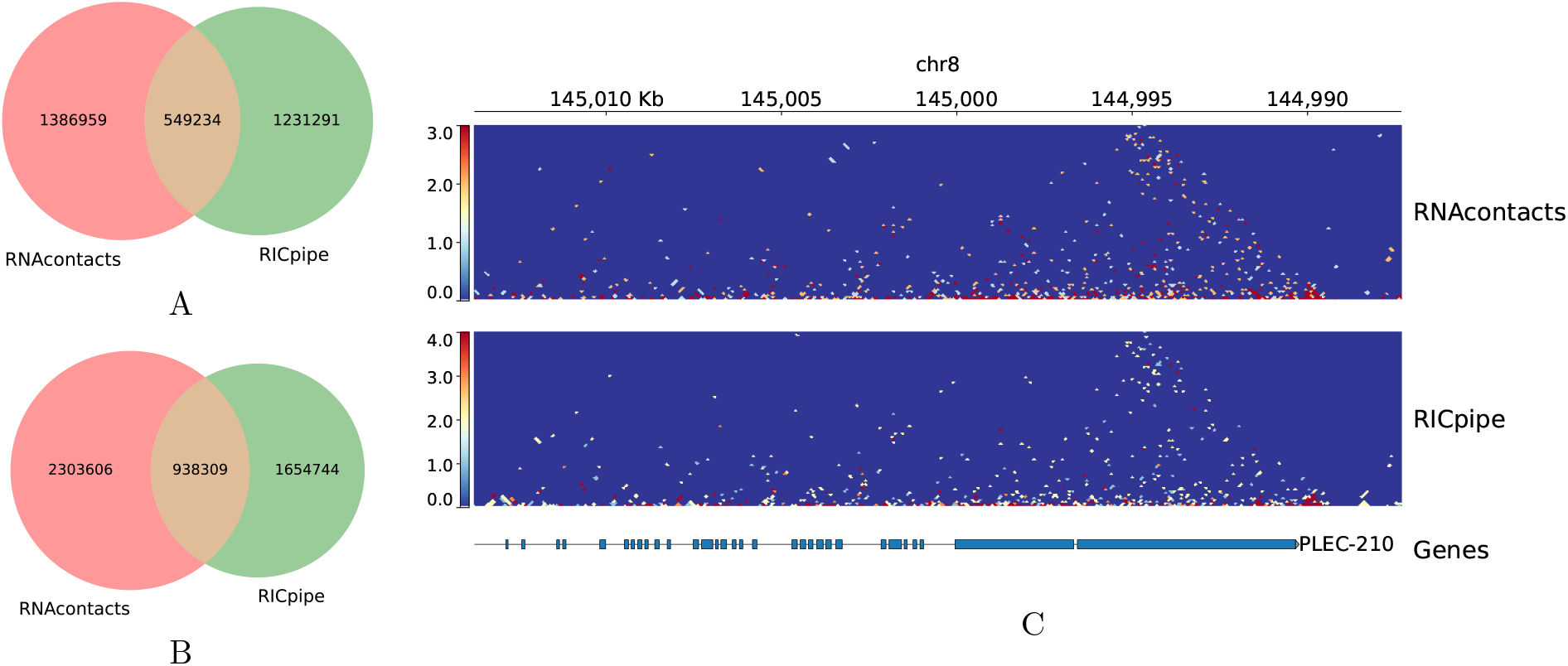
RNAcontacts vs. RICpipe. **(A)** Venn diagram of ligation junctions obtained by *RNAcontacts* and *RICpipe*. **(B)** Same as (A), but weighted by read support. **(C)** Contact maps for the *PLEC-210* gene obtained by *RNAcontacts* (top) and *RICpipe* (bottom).

Additionally, we checked the performance of *RNAcontacts* on the RIC-seq data in the HeLa cell line with and without the first pass. Towards this goal, we ran the second pass of *RNAcontacts* while supplying splice junctions that are annotated in GENCODE [19] as opposed to splice junctions that were inferred for the HeLa cell line on the first pass. As a result, approximately 1% of spurious ligation junctions were obtained, ones that actually correspond to the endogenous splice junctions in HeLa. We also found that 16,809 out of ∼3.5 million ligation junctions identified by *RICpipe* could be attributed to spliceosomal introns. While the number of such ligation junctions is not large, they are supported by a considerable fraction (*>* 30%) of reads. We conclude that two-pass method provides higher specificity (lower false positive rate) towards RNA-RNA contacts, especially in the conditions when the expressed transcriptome differs significantly from the annotation.

## 3 Discussion

In this work, we presented a conceptual solution to the problem of mapping short reads with two distance split types, which are characteristic to RNA proximity ligation assays, using STAR aligner. However, the approach is not limited to STAR, and any other spliced aligner program can be used instead [17]. We demonstrated that endogenous splice junctions indeed constitute a large part of split read alignments in RIC-seq data, and that *RNAcontacts* allows for more sensitive detection of split reads aligning to ligation junctions than *RIC-pipe*. The implementation of *RNAcontacts* in a reproducible and scalable workflow management system *Snakemake* allows fast and uniform processing of multiple datasets like RIC-seq.

The nature of RNA proximity ligation data is similar to that of Hi-C, yet it has important distinctions related to the resolution. While for Hi-C, it is a common practice to average chromatin contacts at kilobase or megabase scale, the assessment of RNA-RNA contacts using proximity ligation intrinsically targets single-nucleotide level. At that, the read support by RIC-seq in most naturally-occurring contacts, e.g, one that is mediated by RNA structure in the human *SF1* gene [20] is very weak (see example in Figure 3). We observed that most RIC-seq contacts *in cis* and *in trans* were supported by only one read. This raises an important question of how to evaluate the statistical significance of the contacting clusters. This question should be addressed in future studies fostered by larger amounts of data. The authors anticipate that many more RNA proximity ligation datasets similar to RIC-seq will soon become available, for which *RNAcontacts* pipeline is readily available.

**Figure 3:**
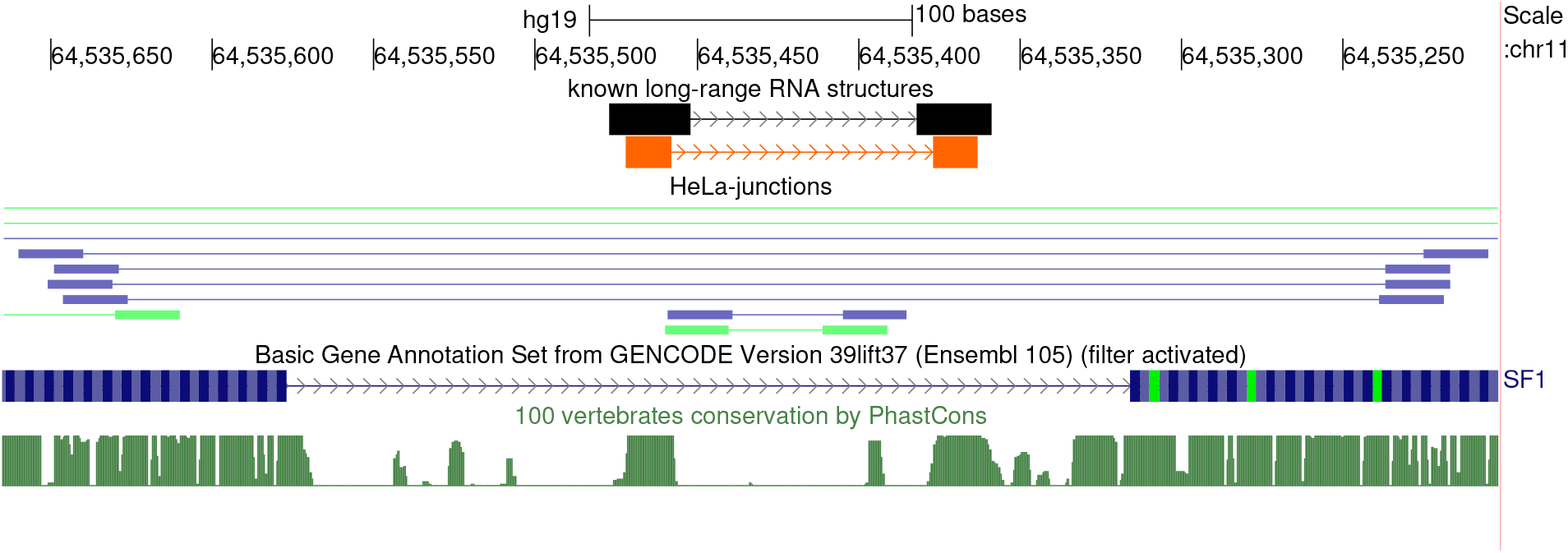
Ligation junctions supporting RNA structure in the human *SF1* gene [20]. The complementary strands are shown in orange. Ligation junctions in HeLa cell line are shown under HeLa junctions track. The reads from the two bioreplicates are shown in blue and green.

## 4 Conclusions

*RNAcontacts* implements a generic pipeline for the analysis of RNA-RNA contacts that accounts for multiple split types specific for RNA proximity ligation methods. This software was initially designed to be used for the RIC-seq protocol, however its scope extends to any method involving proximity ligation, in which one of the interacting partners is RNA.

### 4.1 Methods

#### 4.2 Genomes and annotations

February 2009 (hg19) assembly of the human genome and GENCODE transcript annotation v34lift37 were downloaded from Genome Reference Consortium [21] and GENCODE website [19], respectively. Intron coordinates were taken from STAR output (see below).

#### 4.3 High-throughput sequencing data

Two bioreplicates of rRNA depleted RIC-seq data (GSM3629915 and GSM3629916) in the HeLa cell line [14] were downloaded from the Gene Expression Omnibus under the accession number GSE127188 in FASTQ format. The matched control set of RNA-seq data in the HeLa cell line were downloaded from the ENCODE consortium under the accession numbers ENCLB555ASI and ENCLB555ASJ. On the first pass, the RNA-seq data were mapped to the human genome using STAR aligner version 2.7.3a in paired-end mode with the following additional settings: `--runMode alignReads --outSAMtype BAM SortedByCoordinate--chimOutType Junctions`. On the second pass, the RIC-seq data were mapped to the human genome using the same version of STAR aligner with the following additional settings: *--chimSegmentMin 15 --chimJunctionOverhangMin 15 --chimScoreJunctionNonGTAG - 1 --scoreGapNoncan -1 --scoreGapATAC -1 --scoreGapGCAG -1 –chimSegmentReadGapMax 3 --outFilterMatchNminOverLread 0*.*5 --outFilterScoreMinOverLread 0*.*5*. The parameter *– chimSegmentReadGapMax 3* is introduced to account for mappability of the additional biotinylated cytosine residue in RIC-seq [14]. The penalty score is reduced to −1 for all non-canonical splice junctions on the second pass.

#### 4.4 Pipeline implementation

*RIC-contacts* is implemented in the popular workflow management system *Snakemake* [22] and is freely available through GitHub [23]. The input data files are provided through a configuration file in yaml format, which also contains STAR settings and additional parameters that control the minimum distance between two ligation points in a cluster and the cutoff on the distance between neo-junctions to be visualized through UCSC Genome Browser [24]. Neojunctions were extracted from BAM files using custom Perl script (textttneo.pl in *RNAcontacts* repository) and *samtools* package v1.14 [25]. *bedops* package v2.4.41 was used to cluster ligation points [26]. The number of supporting reads was computed using *bedtools* package v2.29.0 [27].

#### 4.5 Visualization

For contact map visualization we convert the contact lists to ‘cool’ format using ‘cooler’ suite v0.8.11 with 100bp resolution and visualize the maps with pygenometracks v3.7. Split read visualization is performed with IGV v2.11.2 and the UCSC Genome Browser [24]. By default, only co-linear contacts that span not more than 50,000 nts are visualized through a UCSC Genome Browser track hub (see also the manual [23]).

## 5 Availability of source code and requirements

- Project name: RNAcontacts
- Project home page: https://github.com/smargasyuk/RNAcontacts
- Operating system(s): GNU/Linux
- Programming language: Snakemake
- Other requirements: Python 3.7 or higher, Snakemake 7.12.0 or higher, conda 4.11.0 or higher, mamba 0.19.0 or higher
- License: GNU GPL v3

## 6 Availability of supporting data and materials

The data set supporting the results of this article is available in the Zenodo repository (https://zenodo.org/record/7027475) [28].

## 7 Declarations

### 7.1

#### List of abbreviations

EEJ: Exon-exon junction
RIC-seq: RNA in situ conformation sequencing
RBP: RNA-binding protein

### 7.2 Consent for publication

Not applicable.

### 7.5 Author’s Contributions

SM and DP designed the study. SM implemented the computational pipeline. MV, GL, CC, and DP analyzed the data. SM and DP wrote the first draft of the manuscript. All authors edited the final version of the manuscript.

## 8 Acknowledgements

Authors thank Timofei Ivanov for additionally testing the software.

