## Supplementary information for "RNAcontacts, a pipeline for predicting contacts from RNA proximity ligation assays"

September 6, 2022

### List of Figures

### List of Tables

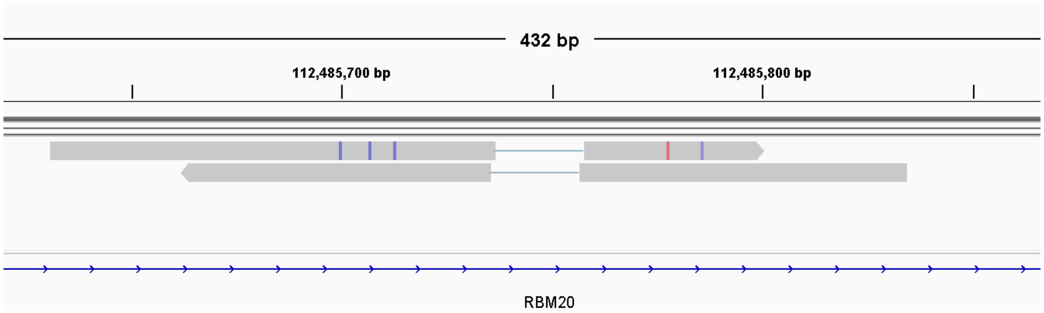


**Figure S1:** An example demonstrating that the variability in gap positions can arise even when mapping read mates that overlap the same ligation point, which were sequenced from opposite strands.

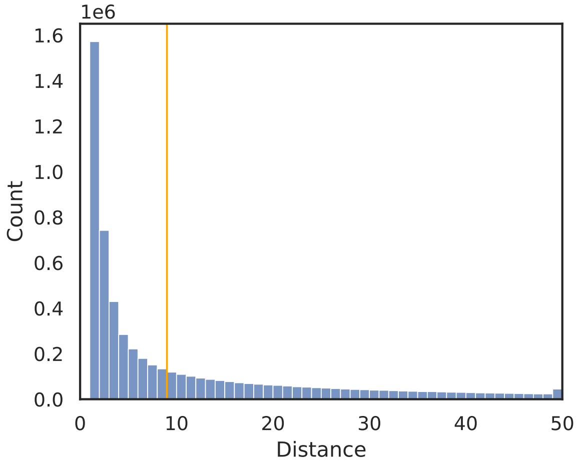


**Figure S2:** The distribution of distances between two consecutive ligation points obtained after the second pass of `RNAcontacts` pipeline. The vertical line represents the median of the distribution.

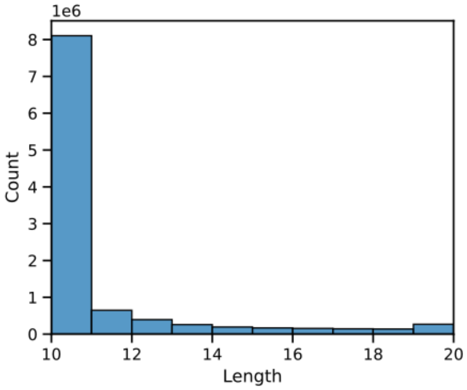


**Figure S3:** The distribution of cluster lengths with the clustering distance $\delta = 10$.

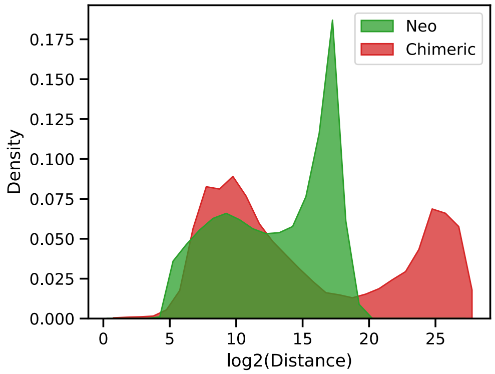


**A** (by positions)

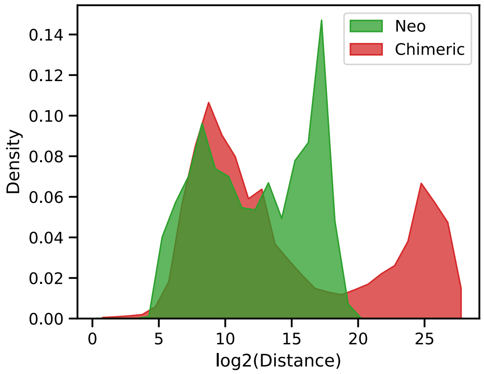


**B** (by number of reads)

**Figure S4:** The distribution of distances between contacting points for neo-junctions and chimeric splits, in terms of the number of split positions (A) and weighted by the number of supporting reads (B).

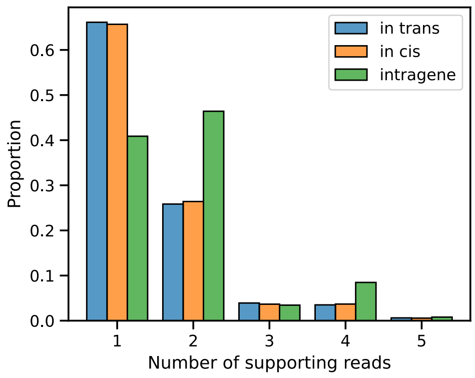


**Figure S5:** The distribution of the number of reads supporting the contact.

| Rep | Mate | Total | Mapped | Unique | Multimap | % Mapped | % Unique |
|---|---|---|---|---|---|---|---|
| 1 | 1 | 56,329,082 | 53,567,065 | 38,346,521 | 15,220,544 | 95.10% | 71.59% |
| 1 | 2 | 56,329,082 | 52,735,572 | 37,196,880 | 15,538,692 | 93.62% | 70.53% |
| 2 | 1 | 55,853,750 | 53,039,695 | 38,924,672 | 14,115,023 | 94.96% | 73.39% |
| 2 | 2 | 55,853,750 | 52,181,644 | 37,764,455 | 14,417,189 | 93.43% | 72.37% |
| | Total | 224,365,664 | 211,523,976 | 152,232,528 | 59,291,448 | 94.28% | 71.97% |

**Table S1:** Read mapping statistics for the HeLa cell line. Columns (left to right): bioreplicate number, mate (read1/read2), total number of reads, the number of mapped reads, the number of uniquely mapped reads, the number of multimaps, % reads that were mapped, % of reads that were mapped uniquely among reads that were mapped.
